# The natural HLA-Peptidome of Sezary Syndrome: uncovering antigens for T cell-based immunotherapy

**DOI:** 10.1101/2024.12.25.630030

**Authors:** Lydon Wainaina Nyambura, Kathrin Textoris-Taube, Peter Walden

## Abstract

Sezary syndrome (SS) is a rare and aggressive form of cutaneous T-cell lymphoma, that has a poor prognosis, with a median overall survival time of less than 3 years. Despite advances in its treatment, SS is a challenge to manage, often characterized by high rates of relapse and limited response to therapy in many patients. The main challenge for treatment, including vaccine development is its heterogeneity. In its molecular and genetic characteristics, clinical presentation, disease progression, and treatment response. Understanding the SS heterogeneity at the omics level is vital in developing T-cell mediated immunotherapeutic. In this study naturally presented HLA-I peptides were isolated from SS patients leukophoresis samples and analyzed using high performance liquid chromatography coupled to tandem mass spectrometry (LC-MS/MS), and source proteins evaluated for immunotherapy application. The total number of HLA-I–restricted peptides and source proteins identified in SS leukophoresis patient samples was about the same, and they were heterogeneous and individualized. Only a small fraction of HLA-I peptides and source proteins was found to be shared between and among the patients. Peptide lengths were dominated by nanopeptides, with preferred processing by chymotrypsin. The source proteins were predominantly from the cytoplasm, and primarily involved in biosynthesis and regulation. Furthermore, the HLA-I peptides were presented from top 20 genes with somatic mutations in SS that include NCOR1, TRRAP, JAK3, PLCG1, TP53 and STAT3 (SS associated antigens-SAAs). These SAAs had; varying mutation types and frequencies-dominated by missense variant, allele-dependent immunogenicity-highest in HLA-A*11:01 and HLA-A*02:01 and lowest for HLA-A*01:01-with TRRAP showing high affinity peptides, low gene expression levels in normal tissue (except STAT3), significant protein interaction network-including JAK3 and STAT3 at primary level. This study findings, contributes to the overall understanding SS HLA-I peptidome landscape, and highlights potential T-cell mediated immunotherapeutic targets.

## Introduction

Sezary syndrome (SS) is a rare and aggressive form of cutaneous T-cell lymphoma, characterized by the proliferation and accumulation of malignant T cells in the skin (Alaggio & Amador, 2022; Willemze et al., 2019). SS typically presents with generalized erythroderma, intense pruritus, and the presence of atypical Sezary cells in the peripheral blood (Willemze et al., 2019). The disease primarily affects older individuals, with a higher incidence in males (Agar et al., 2010). Unfortunately, SS has a poor prognosis, with a median overall survival time of less than 3 years (Agar et al., 2010; Scarisbrick et al., 2015). Therapeutic management of SS involves a multimodal approach, combining skin-directed therapies (topical corticosteroids, phototherapy, and local radiotherapy), systemic treatments such as retinoid (Jawed et al., 2014; Querfeld et al., 2019; Stuver & Geller, 2023). Chemotherapy, and targeted agents, are utilized for more advanced stages or refractory disease (Jawed et al., 2014; Querfeld et al., 2019; Stuver & Geller, 2023). Immunotherapies, including immune checkpoint inhibitors and adoptive T-cell therapy, show promise in enhancing anti-tumor immune responses (Roccuzzo et al., 2021). Despite these advances in treatment, SS remains challenging to manage, often characterized by high rates of relapse and limited response to therapy in many patient (Larocca & Kupper, 2019).

The main challenge for SS treatment, including vaccine development is its heterogeneity (Buus & Willerslev-Olsen, 2018; Litvinov et al., 2017; Stuver & Geller, 2023; Vergnolle et al., 2022). In its clinical presentation, disease progression, and treatment response. Including, molecular and genetic characteristics among patients, leading to distinct patterns of gene expression and signalling pathways. Understanding the SS heterogeneity at the omics level is vital in developing T-cell mediated immunotherapeutic approaches, such as peptide based vaccines.

HLA peptidomics refers to the study of the peptide repertoire presented by human leukocyte antigen (HLA) molecules (Bassani-Sternberg et al., 2016; Caron et al., 2015; Hellinger & Sigurdsson, 2023; Kalaora et al., 2016; Shapiro & Bassani-Sternberg, 2023; Vaughan et al., 2017). These peptides are processed from intracellular proteins primarily by the proteasome and presented to the cell surface by HLA for T-cells recognition. HLA peptidomics allows for understanding the mechanism of antigen processing and presentation in cancer cells. Including identification and characterization of peptides from tumour associated antigens (TAAs) and neoepitopes presented by HLA molecules on cancer cells. Overall, HLA peptidomics in cancer represents a powerful tool for understanding the mechanism of antigen processing and presentation, highlighting potential tumour targets, for T-cell mediated immunotherapies.

HLA peptidomics study for SS, is warranted to determine how this SS heterogeneity impacts antigen processing and presentation, and identify potential targets for SS T-cell mediated immunotherapeutic vaccine. In this study, immunoaffinity purification of HLA-I peptide complexes from SS clinical leukophoresis patients’ samples was carried out and analyzed by high-performance liquid chromatography coupled to tandem mass spectrometry (LC-MS/MS). These HLA peptidomes were studied and compared in reference to presentation, lengths, subcellular locations, molecular/biological functions, and select SAAs analysed according to their immunogenicity, gene expression profiles, and protein interaction partners.

## Materials and Methods

### Sezary Syndrome patients

The clinical leukephoresis material from 4 SS patients was used with approval by Charité ethics committee (Approval No. EA1/222/14 and EA1/026/14) and written informed consent by the volunteer donors. HLA type was determined by Charité - Universitätsmedizin Berlin, HLA typing laboratory.

### Isolation and purification of MHC I-presented peptides

MHC class I molecules were isolated as described in detail elsewhere (Nyambura et al., 2016). Briefly, cells were lyzed in 20 mM Tris-HCl buffer, pH 7.4, 0.3% CHAPS, 0.2% NP-40, 145 mM NaCl, 1 mM EDTA, 1mM Pefabloc. Lysates were ultracentrifugated for 1h at 100,000 x g and the supernates passaged through a column with monoclonal antibody of irrelevant specificity followed by a column with the HLA-I anti-human monoclonal antibody (W6/32), both coupled to activated CH Sepharose as per manufacturer’s protocol (Amersham Biosciences AB, Uppsala, Sweden). After adsorption of the proteins, the anti-human HLA-I column was washed with the following in descending order; 20mM Tris, 145 mM NaCl, pH 7.4 (TBS), 0.3% CHAPS in TBS, TBS, 0.3% ß-octylglycoside in TBS, TBS, and lastly with ultrapure H_2_O. HLA-peptide complexes were eluted from the column with 0.7 % TFA in ultrapure H_2_O. High molecular weight components were separated from peptides by centrifugal ultrafiltration using a molecular weight cut-off 3 kDa (Centricon, Millipore, Schwalbach, Germany). The filtrates were fractionated on a Smart HPLC system (Amersham Biosciences, Freiburg, Germany) using a reverse phase column µRPC C2/C18, SC2.1/10 (Amersham Biosciences) and an acetonitrile gradient of 5 - 90% of B (solvent B: 0.1 % TFA, 90% of acetonitrile; solvent A: 0.1% TFA in ultrapure H_2_O). The fractions obtained were lyophilized and re-dissolved in 0.1% TFA 2% acetonitrile for LC-MS/MS.

### LC-MSMS analysis of HLA ligands

The peptide fractionates were analyzed by reversed-phase LC (Ultimate 3000 RSLCnano) coupled on-line with Q Exactive Plus MS (both, ThermoFisher Scientific). Fractionated peptides were trapped on a C18 precolumn at 20 μL/min (2% acetonitrile, 0.1% TFA) for 4 min. Subsequently, peptides were separated at a flow rate of 300nl/min onto a 75-μm × 25cm PepMap nano-HPLC column with a gradient of 3–30% of 80% acetonitrile and 0.1% FA acid in ultrapure H_2_O over 90 min. Eluted peptides were nanospray-ionized and fragmented based on the ten most intense precursor ions signals, with a 20 sec dynamic exclusion time to avoid repeated fragmentation.

### Data processing and analysis

The MS and MS/MS spectra were processed via Data Analysis (vers. 3.4) and Bio-tools (vers. 3.1) software (Bruker Daltonics). The local MASCOT server (vers. 2.2), utilising the Swissprot databank (vers. 56.3) for human proteins (20,408 reviewed entries), was used to identify the peptides. The precursor mass tolerance was 5 ppm, and 10 ppm for MS/MS, and methionine oxidation as possible variable modification. For each peptide-spectrum match, candidate sequences were validated using a statistical evaluation -10logP, where logP is the logarithm to the base 10 of P (P<0.05) as the absolute probability. Further validation of the identified peptides on the basis of *de novo* sequencing was done using the Sequit software (Demine & Walden, 2004), and by manual inspection of the peptide-spectrum. The protein sequence, protein ID and gene symbol for proteomic data analyses were extracted from Uniprot database (“UniProt: The Universal Protein Knowledgebase in 2023.,” 2023) . The human protein reference database (Mishra et al., 2006), was used to classify proteins according to their subcellular localizations and biological function.

### Somatic mutations

To determine the top 20 genes with somatic mutation in SS, the Catalogue Of Somatic Mutations In Cancer (COSMIC) was used https://cancer.sanger.ac.uk/cosmic (Sondka et al., 2024). The main search was set to SS, using the following browser filters; Tissue selection (Hematopoietic and lymphoid), Sub-tissue selection (All), Histology selection (lymphoid neoplasm) and Sub-histology selection (mycosis fungoides-sezary syndrome). In addition, the type of somatic mutation of the SAAs (The top 20 genes with somatic mutation in SS as per the COSMIC database and those whose HLA-I peptides were presented by all the 4 patients or at least 3 of the four patients (Rom, IrK, FrA and Seo), was also determined by COSMIC.

### Sezary syndrome associated antigens (SAAs) immunogenicity

To determine the immunogenicity of the SAAs, NetMHCpan 4.1 BA in the IEDB (http://www.iedb.org) was used (Vita et al., 2019). The immunogenicity was determined for HLA-A*01:01, HLA-A*02:01, HLA-A*11:01, HLA-A*24:02, HLA-C*06:02, HLA-C*07:01 and HLA-C*07:02, individually and all combined. These alleles together are expressed in 90% of the population (Gonzalez-Galarza et al., 2020). IC50(500) nM binding affinity threshold was used as threshold for immunogenicity, and immunogenicity scores were presented as 1/IC50(500) nM.

### SAAs gene expression in major normal human tissues, and protein interaction partners

To determine the gene expression profiles of the SAAs in normal major human tissues, the genecard database was used https://www.genecards.org/ (Stelzer et al., 2016). With a low gene expression intensity cut-off of 10%. The SAAs protein interaction partners was determined using STRING vers.11.5; a database of known and predicted protein-protein interactions (Szklarczyk et al., 2019). STRING was used to determine known SAAs protein interaction partners, experimentally determined from various biochemical, biophysical and genetic techniques. A medium interaction score of 0.400 was applied with a cut-off of 10 interaction partners at primary level against *Homo sapiens*.

## Results

### Naturally presented HLA I ligands of Sezary syndrome patients

The SS leukophoresis patient samples (Rom, IrK, FrA and Seo) were lysed, and MHC class I molecules isolated by affinity chromatography, and peptides extracted from the MHC molecules analyzed by LC-MS/MS. The sequences of a total of 7668, 8492, 8123 and 4854 HLA class I-bound peptides were identified from 3003, 4027, 3865 and 2390 source proteins in Rom, IrK, FrA and Seo respectively **(Fig. 1A)**. 6086, 7512, 6905 and 3441 HLA class I-bound peptides were unique in Rom, IrK, FrA and Seo, respectively **(Fig. 1A)**. The shared peptides between the patients ranged from 146 (0.6%) - 748 (2.8%), and well below 1% among the patients. Only 69 (0.3%) HLA I peptides sequences were found to be shared among all the 4 patients based only on peptide sequences, and precursor peptide mass signals and retention time respectively **(Fig. 1B)**. The HLA expression was HLA-A2, HLA-A24, HLA-B7 and HLA-B8, HLA-C6 and HLA-C7 in Seo, HLA-A1, HLA-A33, HLA-B8, HLA-B14, HLA-C7 and HLA-C8 in Rom, while the HLA expression in IrK and FrA was undetermined. The shared proteins between the patients ranged from 187 (2.6%) to 652 (9.1%), and only 627 (8.7%) among the 4 patients. **(Fig. 1C)**

**Figure 1:**
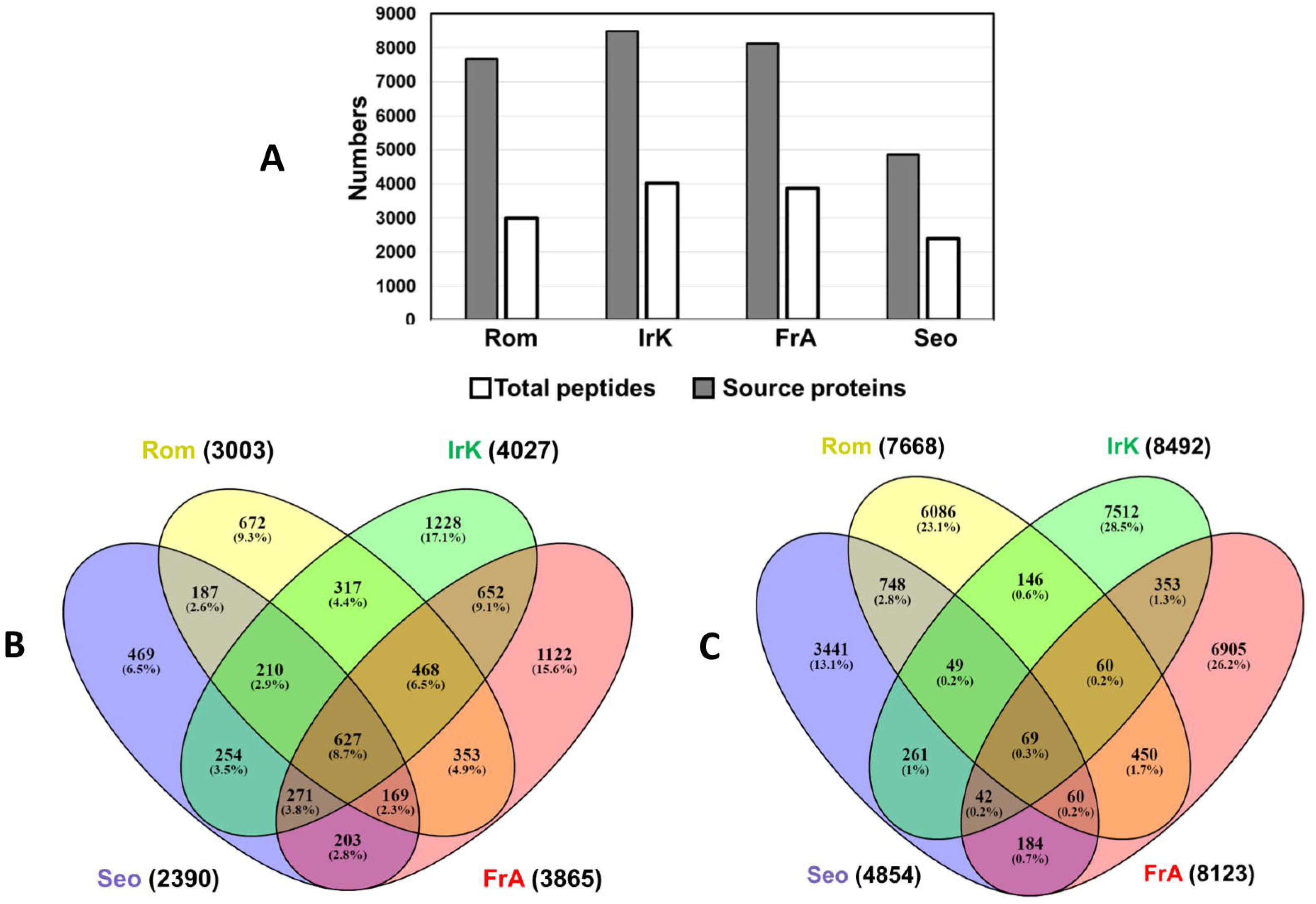
Naturally presented HLA class I ligands and Source proteins of Sezary syndrome leukapharesis patients samples. The leukapharesis samples were lysed, MHC I molecules were isolated by affinity chromatography, and peptides extracted from the MHC molecules were analyzed by LC-MS/MS. **A)** Number of naturally presented HLA class I ligands and Source proteins. **B)** Shared-Naturally presented HLA class I ligands**. C)** Shared-Naturally presented HLA class I ligands source proteins.

### MHC I-bound peptide lengths and C-Terminus processing

The MHC I-bound peptide lengths in all patients were dominated by nanopeptides constituting 29.0% -55.1% of all the identified peptides. While decapeptides were the second dominant, constituting 8.0% -14.8%. Undecapeptides and above, on the other hand were less than 11% in all patients **(Fig. 2)**. The C-terminus peptide processing by the proteasome based on the peptide numbers and percentages was Chymotrypsin>Trypsin>Caspase in all patients **(Fig. 3)**.

**Figure 2:**
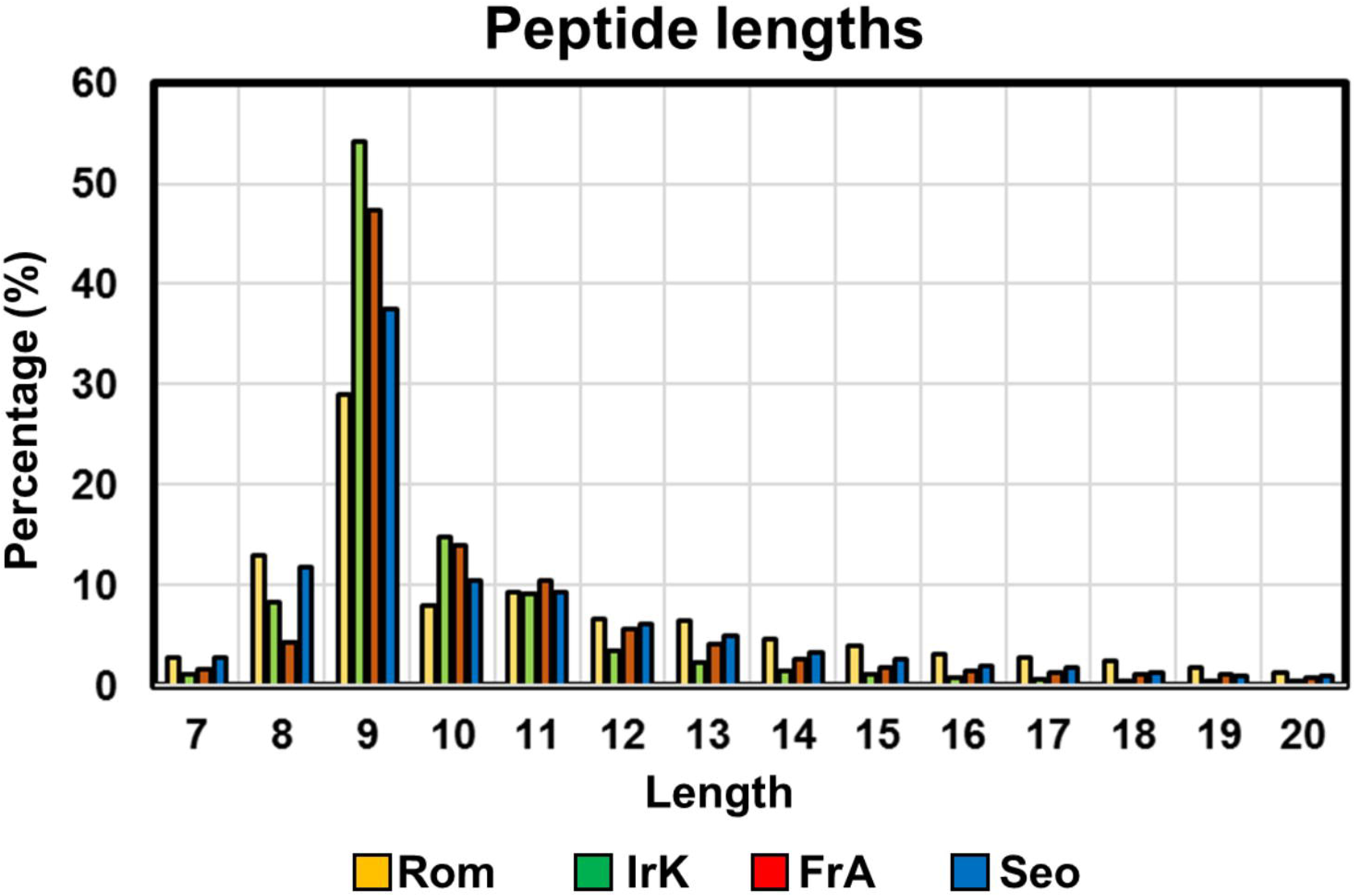
MHC Class I -peptide lengths in Sezary syndrome Leukapheresis patients’ samples.

**Figure 3:**
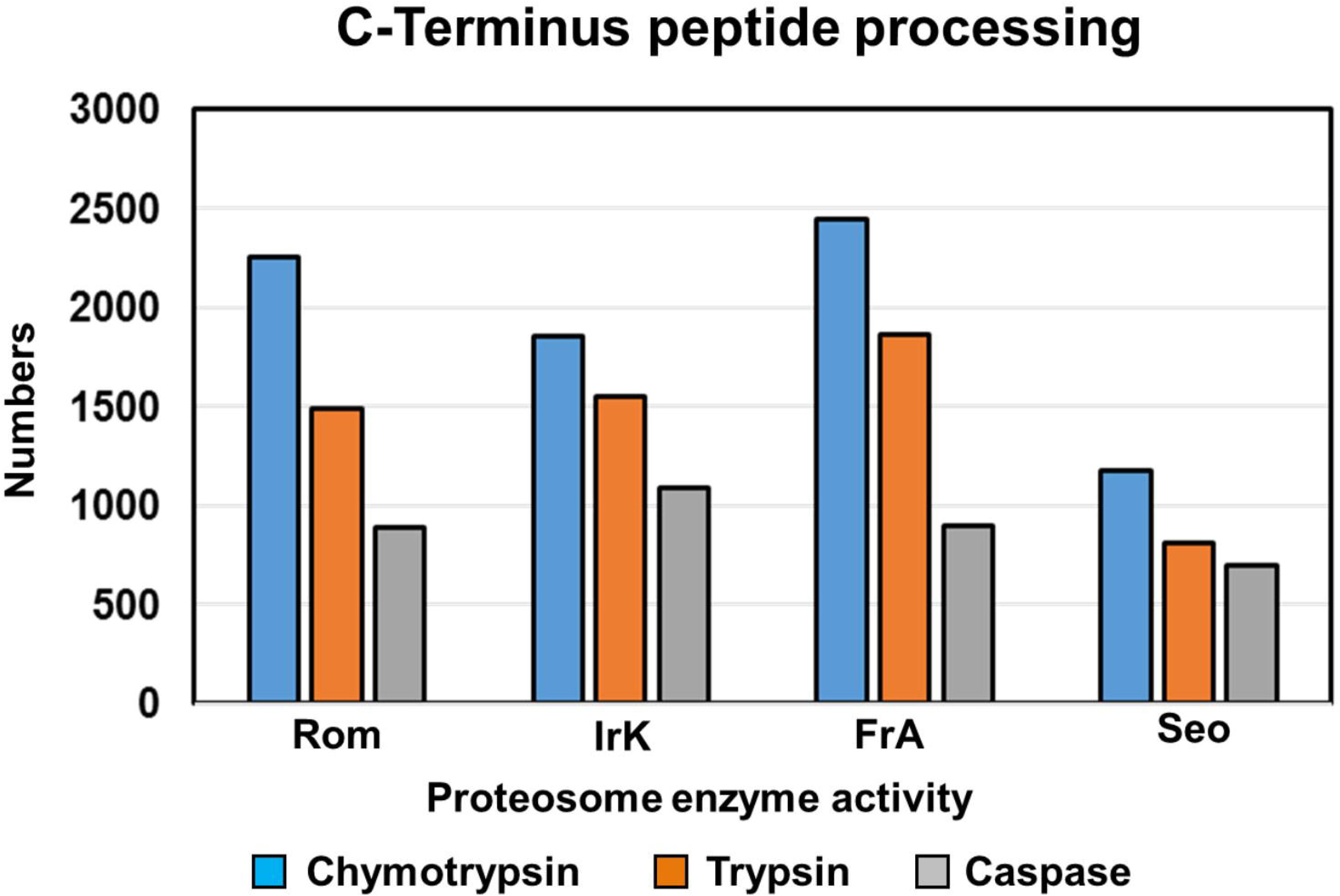
MHC Class I –peptides C-Terminus processing by the proteasome.

### Sub-cellular locations and molecular functions of source proteins

The human protein reference database was used to assign the subcellular location of the source proteins of the HLA I-bound peptides from Rom, IrK, FrA and Seo. The cytoplasm part, intracellular membrane-bounded organelle and cytosol were the dominant subcellular locations of the source proteins **(Fig. 4)**. Together, they accounted for 29.6 %, 28.8 %, 27.1 % and 24.7% of all the source proteins in Rom, IrK, FrA and Seo, respectively. Approximately half of this percentage was solely from the cytoplasmic part, which accounted for more than 14% of all the source proteins. Source proteins from the cell junction was 10.3% in Seo, considerable high compared to those in Rom, Irk and FrA which were 3.9 %, 3.9% and 3.3% respectively. Source protein subcellular locations were lowest in condensed chromosome, endoplasmic reticulum lumen and intracellular organelle lumen in all patients with values ranging from 0% - 0.32% (data not shown). The source proteins were from diverse subcellular locations in all patients (**Fig. 4)**. The source proteins were further evaluated for their biological/molecular functions, again using the human protein reference database (Mishra et al., 2006). Although source proteins possessed multiple biological/molecular functions, a vast majority were involved in biosynthetic process (14.7%, 13.4%, 12.0%, 11.6%), biological regulation (8.9%, 10.0%, 10.5%, 11.1%), catabolic process (8.1%, 5.3%, 4.8%, 5.6%) and anatomical structure development (5.3%, 4.1%, 4.6%, 4.4%) in Rom, IrK, FrA and Seo respectively. **(Fig. 5)**. They were also those involved in immune response, cell death and apoptosis, but were below 2% in all patients.

**Figure 4:**
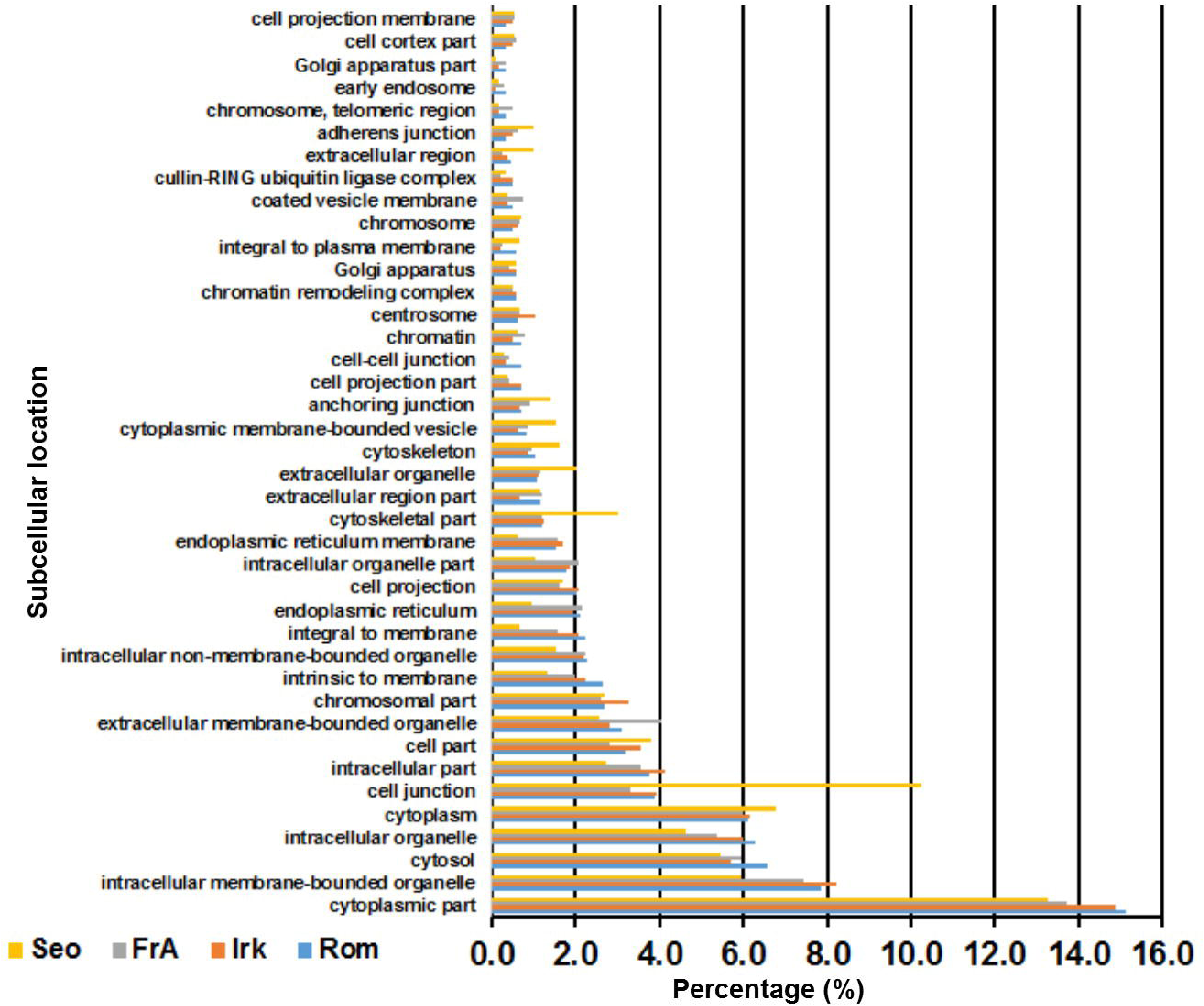
Subcellular location of the source proteins. The subcellular locations of the source proteins of the HLA class I-bound peptides from Sezary syndrome Leukapheresis patients’ samples identified by mass spectrometry, were assigned using the Human Protein Reference Database.

**Figure 5:**
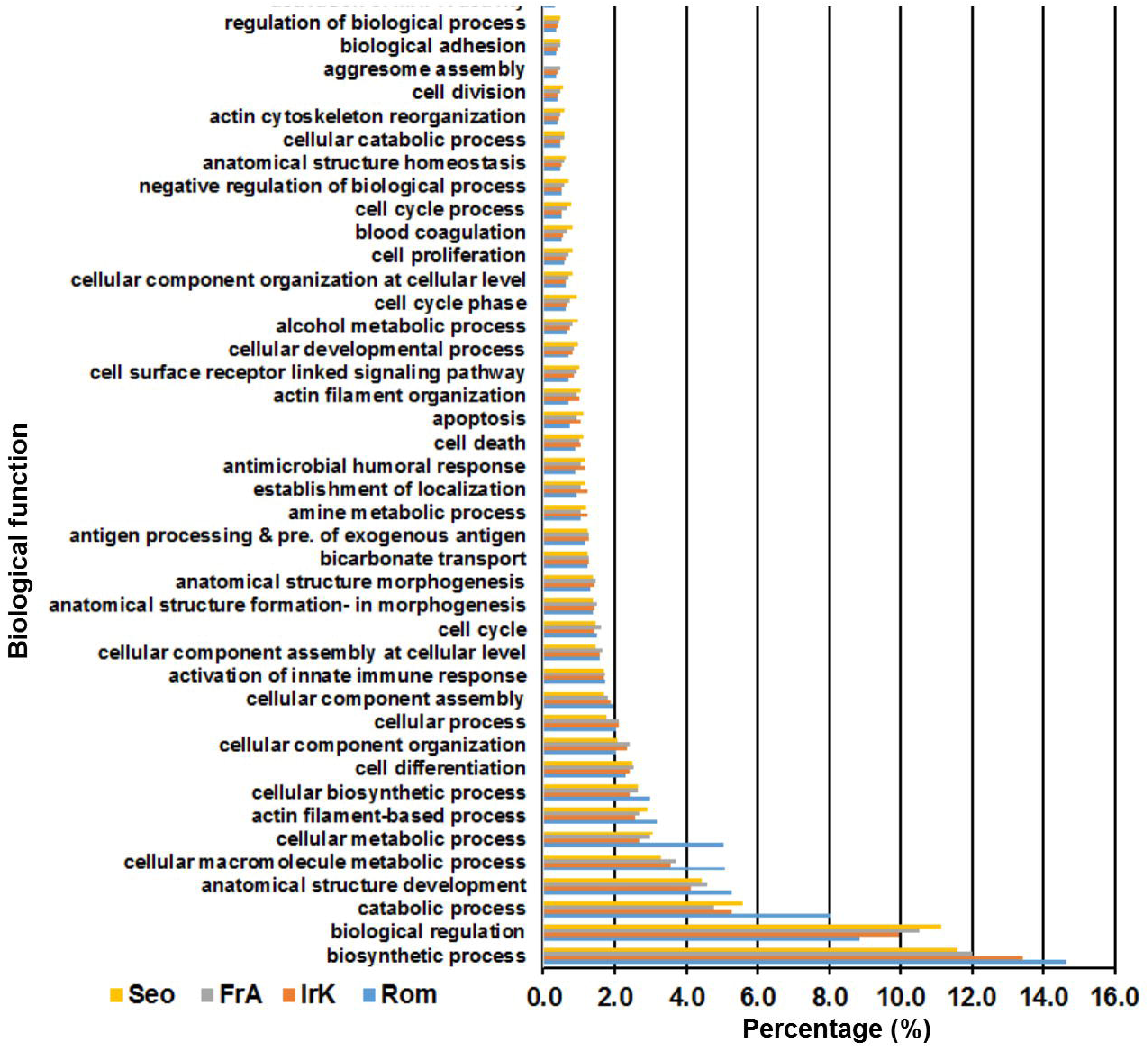
Biological/molecular functions of the source proteins. The biological and molecular functions of the source proteins of the HLA class I-bound peptides from Sezary syndrome Leukapheresis patients’ samples, were assigned using the Human Protein Reference Database.

### Somatic mutations in cosmic

To determine the top 20 genes with somatic mutation in SS the catalogue of somatic mutations in cancer (COSMIC) was used. The top 20 genes with somatic mutation as per the COSMIC, search filtered as follows; tissue (hematopoietic and lymphoid), sub-tissue (All), histology (lymphoid neoplasm) and sub-histology (mycosis fungoides-sezary syndrome) are Fat Atypical Cadherin 4 (FAT4), Fat Atypical Cadherin 1 (FAT1), Caspase Recruitment Domain Family Member 11 (CARD11), Nuclear Receptor Corepressor 1 (NCOR1), Phospholipase C Gamma 1 (PLCG1), Tumor Protein P53 (TP53), Glutamate Ionotropic Receptor NMDA Type Subunit 2A (GRIN2A), AT-Rich Interaction Domain 1A (ARID1A), LDL Receptor Related Protein 1B (LRP1B), phosphatidylinositol-3,4,5-trisphosphate dependent rac exchange factor 2 (PREX2), Protection of Telomeres 1 (POT1), Tet Methylcytosine Dioxygenase 2 (TET2), DNA Methyltransferase 3 Alpha (DNMT3A), Protein Tyrosine Phosphatase Receptor Type K (PTPRK), Erb-B2 Receptor Tyrosine Kinase 4 (ERBB4), Signal Transducer and Activator of Transcription 5B (STAT5B), Transformation/Transcription Domain Associated Protein (TRRAP), Janus Kinase 3 (JAK3), Platelet-Derived Growth Factor Receptor Alpha (PDGFRA) and Signal Transducer and Activator of Transcription 3 (STAT3) with mutation frequencies of 16%, 15%, 14%, 14%, 14%, 13%, 11%, 10%, 10%, 10%, 8%, 8%,7%,7%, 6%, 5%, 5%, 4%, 4%, and 4% respectively, from the all the tumors samples **(Fig. 6A)** . HLA-I peptides from NCOR1, TRRAP and JAK3 genes were presented by all the four patients Rom, IrK, FrA and Seo. PLCG1, TP53 and STAT3 by at least 3 of the 4 patients, FAT1 and LRP1B by at least 2 of the 4 patients, FAT4, TET2 and PTPRK by at least 1 of the 4 patients. No HLA-I peptides was presented from FAT4, CARD11, GRIN2A, ARID1A, PREX2, POT1, DNMT3A, ERBB4, STAT5B and PDGFRA **(Fig. 6B)**. The type of somatic in the SAAs (The top 20 genes with somatic mutation in SS as per the COSMIC database and those whose HLA-I peptides were presented by all the 4 patients or at least 3 of the four patients (Rom, IrK, FrA and Seo), that include NCOR1, TRRAP, JAK3, PLCG1, TP53 and STAT3), were predominantly missense variant (a type of substitution in which the nucleotide change results in the replacement of one amino acid with another, a replacement that may alter the function of the protein. With mutation frequencies of 46.7%, 60.0%, 100%, 94.74%, 52.4% and 70% respectively. Nonsense mutations (that occur due to the substitution of a single base pair in a triplet codon, leading to one of three stop codons (UAG, UAA, and UGA). The triplet codon coding for an amino acid is therefore altered to one that prematurely stops mRNA translation and results in a truncated protein. Were found only in NCOR1 and TP53, with 20% and 23% frequencies, respectively (data not shown).

**Figure 6:**
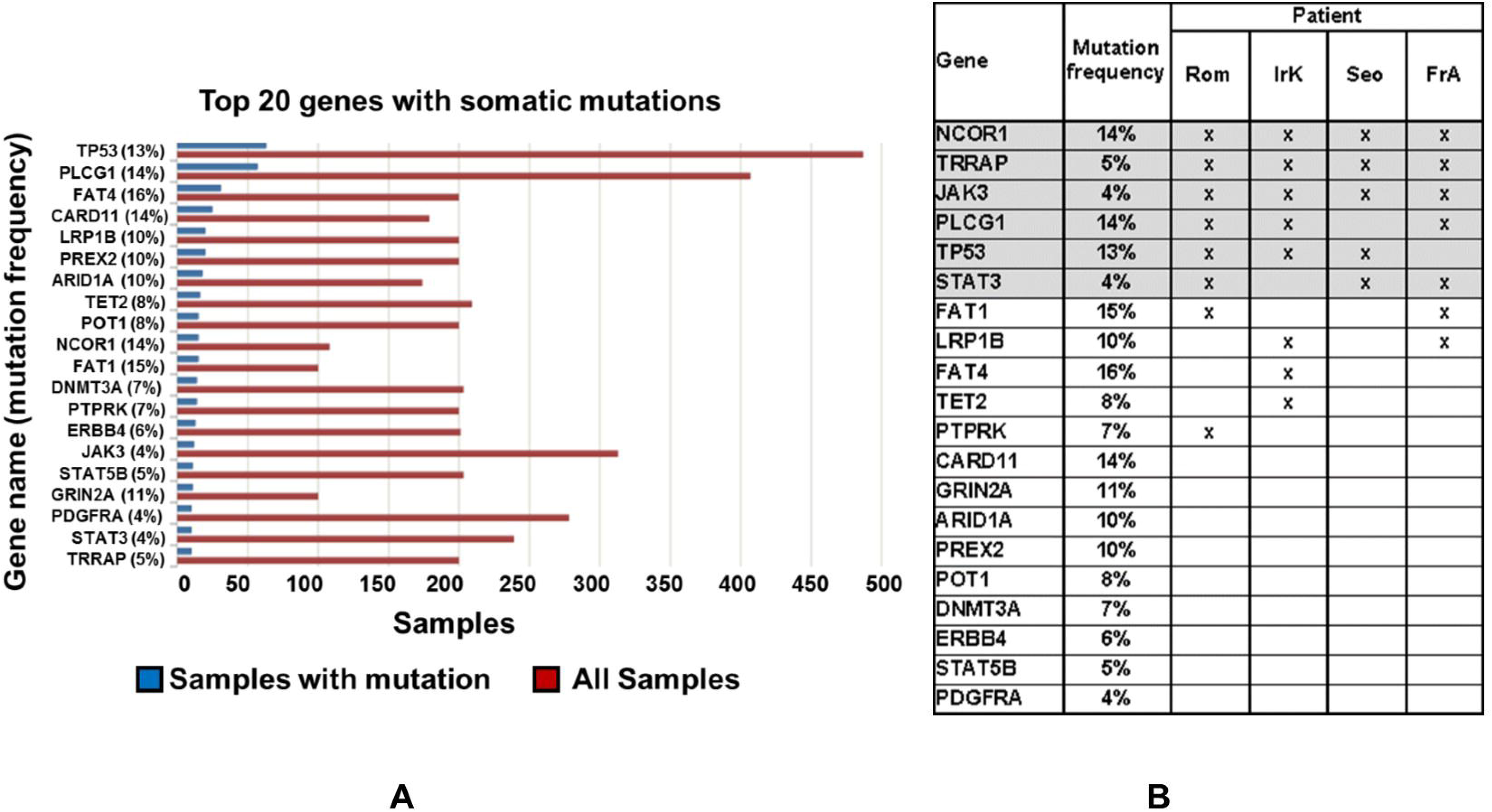
Top 20 genes with somatic mutation in Sezary syndrome as per catalogue of Somatic mutations in Cancer (COSMIC). **A**. COSMIC search set to Sezary syndrome, and filtered by tissue selection (hematopoietic and lymphoid), Sub-tissue selection (All), Histology selection (lymphoid neoplasm) and Sub-histology selection (mycosis fungoides-sezary syndrome). **B**. Frequency of somatic mutation of the SAAs (The top 20 genes with somatic mutation in Sezary syndrome as per the COSMIC database and HLA-I peptides presentation from the SAAs by the patients (Rom, IrK, FrA and Seo).

### Immunogenicity of the LAAs

The immunogenicity of the SAAs (The top 20 genes with somatic mutation in SS as per the COSMIC database and whose HLA-I peptides were presented by all the 4 patients or at least 3 of the four patients (Rom, IrK, FrA and Seo); NCOR1, TRRAP, JAK3, PLCG1, TP53 and STAT3 was determined by reverse immunology. NetMHCpan 4.1 BA in the IEDB was used with a binding affinity threshold of IC_50_ (500 nM), for the most frequent HLA alleles HLA-A*01:01, HLA-A*02:01, HLA-A*11:01, HLA-A*24:02, HLA-C*06:02, HLA-C*07:01 and HLA-C*07:02 that together represent ∼90% of the human population. The immunogenicity score is represented as 1/IC_50_(500 nM) and has a value of 0 to 1 for low to high immunogenicity. The immunogenicity of the SAAs varied depending the HLA allele **(Fig. 7)** and was generally highest for HLA-A*11:01 and HLA-A*02:01and lowest for HLA-A*01:01. The immunogenicity of SAAs for all the alleles was comparable, with median score of less than 0.2 for HLA-A*11:01 and HLA-A*02:01 alleles, and less than 0.4 for all the other alleles, except HLA-A*01:01 which ranged between 0.1 and 0.6. TRRAP had the highest number or peptides within this binding affinity threshold of IC50 (500 nM).

**Figure 7.**
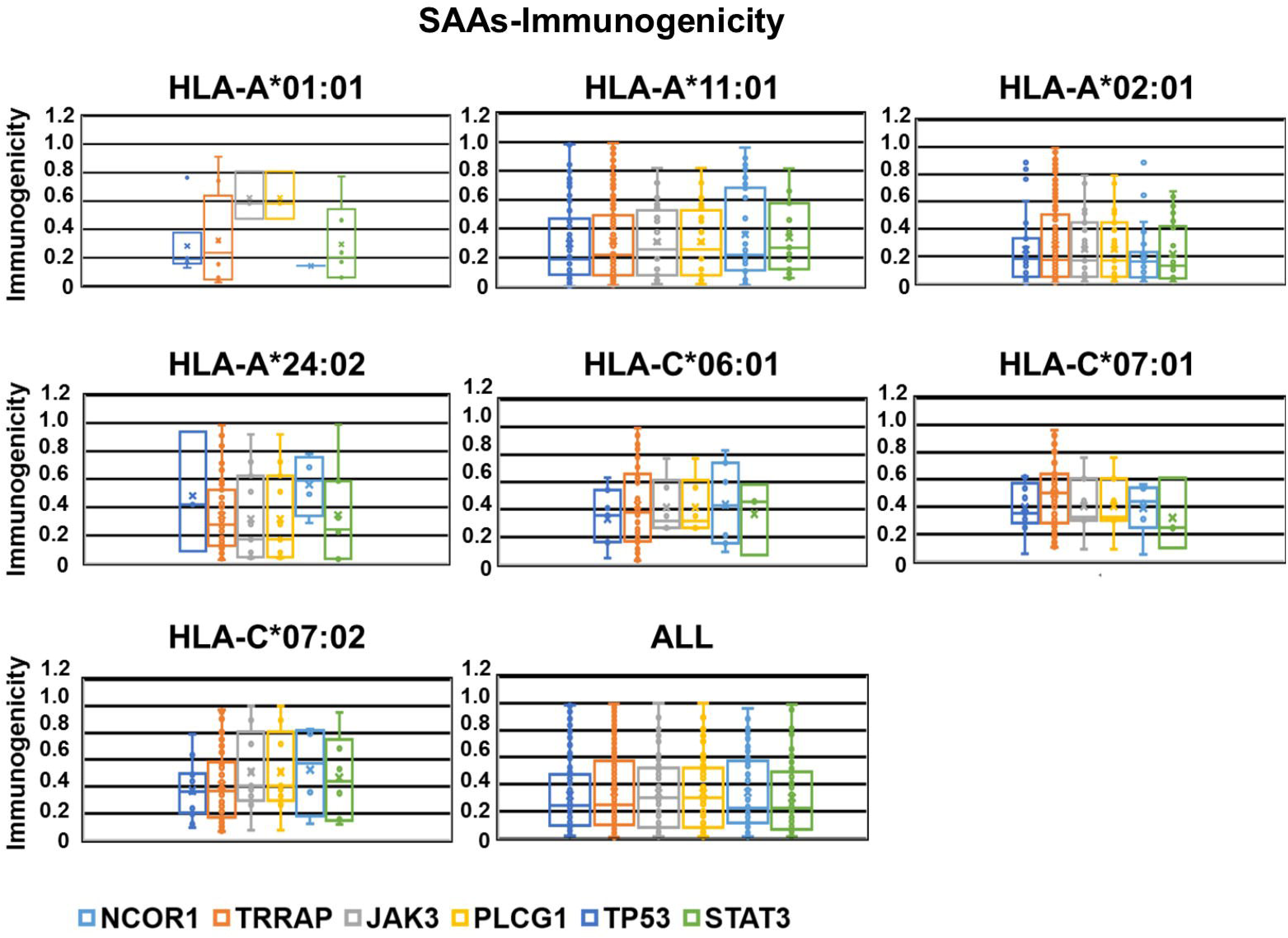
Immunogenicity of the SAAs for the most frequent HLA alleles. HLA-A*01:01, HLA-A*02:01, HLA-A*11:01, HLA-A*24:02, HLA-C*06:02, HLA-C*07:01 and HLA-C*07:02 that together represent more than 90% of the human population. The immunogenicity was determined with NetMHCpan 4.0 in IEDB ^64^ using a binding affinity threshold of IC_50_ (500 nM). The immunogenicity scores are presented as 1/IC_50_ (500 nM) with values of 0 to 1 for low to high immunogenicity.

### SAA gene expression in major normal tissues

The gene expression profiles of the SAAs in major normal human tissues was compared using the genecard database, as detailed in material and methods. All the SAAs were expressed at low levels in all major normal human tissues based on a 10% gene expression intensity cutoff. Only STAT3 were expressed beyond the 10% cutoff. Beyond this cutoff, STAT3 was expressed in whole blood, thymus, adipocyte, lung and prostate, **(Fig. 8)**.

**Figure 8.**
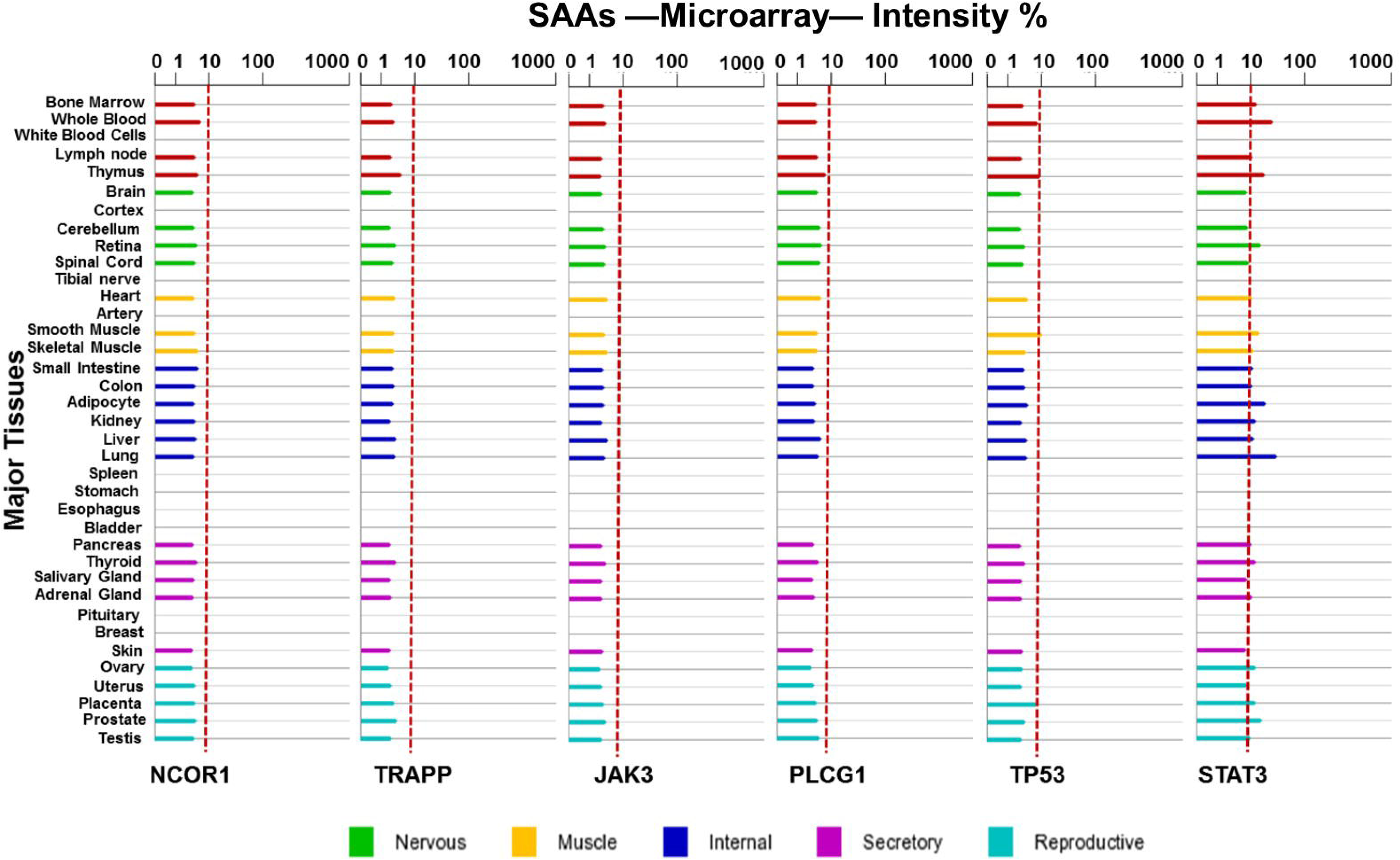
Gene expression profiles of the SAAs in normal human tissue. analyzed and visualized using BioGPS (http://biogps.org/). Low gene expression intensity cutoff of 10% indicated by dotted line.

### Protein interaction partners of the LAAs

High numbers of protein interaction partners and the interaction of SAAs (NCOR1, TRRAP, JAK3, PLCG1, TP53 and STAT3) between and among them may indicate vital roles in SS. The STRING database was used to determine the protein interaction partners of the SAAs, that had been originally identified from experimental data obtained with a variety of biochemical, biophysical and genetic techniques, as detailed in materials and methods. All SAAs (NCOR1, TRRAP, JAK3, PLCG1, TP53 and STAT3) had ten known interaction partners. **(Fig. 9)**. None of the SAAs were found to interact with each other, both at primary (directly via the first shell) (**Fig. 9)** and at secondary level (indirectly via the second shell) (data not shown), except JAK3 and STAT3 that interacted at primary level.

**Figure 9.**
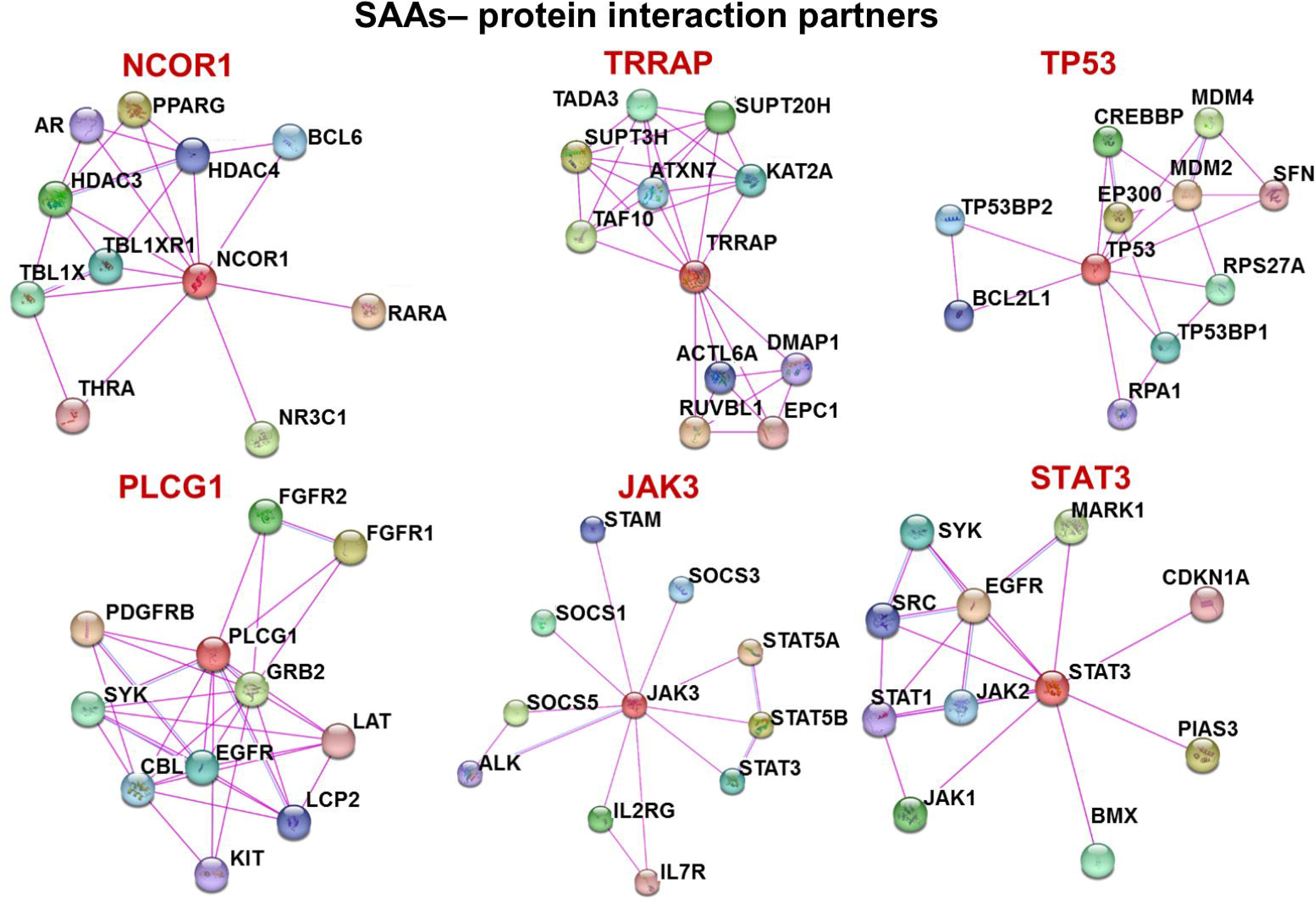
Known protein interaction partners of the SAAs determined using STRING. version 11.0 ^66^ for *Homo sapiens* with a medium score of 0.400 and a cutoff of 10 interaction partners.

## Discussion

The total number of HLA-I–restricted peptides and source proteins identified in SS leukophoresis patient samples was about the same, and they were heterogeneous and individualized **(Fig. 1A. B and C)**. Only a small fraction of HLA-I peptides and source proteins was found to be shared between and among the patients **(Fig. 1B and C)**. The HLA-I–bound peptides were more heterogeneous and individualized than the source proteins, as the number of shared HLA-I peptides was slightly lower compared with those of the source proteins. This has been observed also in other human cell lines and tumor samples such as melanoma (Jarmalavicius et al., 2012; Nyambura et al., 2016; Pritchard et al., 2015; Schellens et al., 2015) and depicts differences in the antigen processing and presentation in SS. In this case, for the HLA-I peptides may also be attributed to the difference in the patient HLA type, though not fully confirmed.

Despite the fact that the HLA-I peptides and the source proteins were heterogeneous and individualized, a number of similarities were observed. First, in all SS leukophoresis patient samples, nanopeptides were the most dominant and constituted between 29.0% to 55.1% of all the identified peptides, followed by decapeptides with 8.0% to 14.8%. **(Fig. 2)**. This nanopeptides dominance has also been observed in other patient tumor samples and cell lines (Fissolo et al., 2009; Jarmalavicius et al., 2012; Nyambura et al., 2016; Pritchard et al., 2015; Schellens et al., 2015), and indicates that the optimum length for MHC I-binding peptides in SS is nine amino acids. Secondly, the proteasome activity in reference to C-terminus peptide processing was in decreasing activity chymotrypsin, trypsin, and lastly caspase, in all patients **(Fig. 3)**.

Similarities were also observed in SS patients’ subcellular locations and molecular functions of HLA-I peptides source proteins. The cytoplasm part, intracellular membrane-bounded organelle and cytosol were the dominant subcellular locations of the source proteins **(Fig. 4)**. Together, they accounted for 24.7% to 29.6 %, with the cytoplasmic part accounting for more than 14% of all the source proteins. This is different to the dominant subcellular locations of the HLA-I peptide source proteins of other human tumor samples and cell lines such as melanoma, multiple sclerosis autopsy samples, B lymphoblastic cell line, tipple-negative breast cancer cell line, leukemia tumor samples and cell lines; where the dominant subcellular locations differed and varied e.g. in melanoma the dominant subcellular locations were the nucleus and the cytoplasm, multiple sclerosis autopsy samples, where cytoplasm and plasma membrane (Brito Baleeiro et al., 2023; Fissolo et al., 2009; Jarmalavicius et al., 2012; Nyambura et al., 2016; Pritchard et al., 2015; Schellens et al., 2015)

Second the source proteins were involved in various molecular functions but especially in biosynthetic process, biological regulation, catabolic process and anatomical structure development, with similar proportion of proteins per molecular function **(Fig. 5)**. This is different to other human tumor samples and cell lines. For instance, in leukemia cell lines MUTZ3 and THP1, the source proteins were involved in cell communication/signal transduction, protein metabolism, and transcription factor activity/regulator activity (Nyambura et al., 2016). While in B lymphoblastic cell line 721.221, the source proteins were dominantly involved in metabolism, cell growth and/or maintenance, cell communication, and stress response (Schellens et al., 2015). In multiple sclerosis autopsy samples, they entailed cellular assembly and organization, nervous system function and development, cellular growth, and proliferation (Fissolo et al., 2009). The similarities in source protein peptide sampling in SS, although unconfirmed, would imply similarities in protein turnover, because protein turnover correlates with source protein peptide sampling (Bassani-Sternberg et al., 2015; Rock et al., 2014). Availability of source proteins involved in immune response, apoptosis, and cell death in all patients, would indicate an active immune activity against the cancer cells by the patients.

Furthermore, the top 20 genes with somatic mutation in SS were, FAT4, FAT1, CARD11, NCOR1, PLCG1, TP53, GRIN2A, ARID1A, LRP1B, PREX2, POT1, TET2, DNMT3A, PTPRK, ERBB4, STAT5B, TRRAP, JAK3, PDGFRA and STAT3, as per COSMIC, with decreasing mutation frequencies of 16% to 4% **(Fig. 6A)**. Out of these, only HLA-I peptides from NCOR1, TRRAP and JAK3 were presented by all the 4 patients, PLCG1, TP53 and STAT33 by 3 of the 4 patients, FAT1 and LRP1B by 2 of the 4 patients, TET2 and PTPRK by 1 of the 4 patients. No HLA-I peptides were presented from FAT4, CARD11, GRIN2A, ARID1A, PREX2, POT1, DNMT3A, ERBB4, STAT5B and PDGFRA **(Fig 6B)**. The missense mutation was the predominant mutation, with mutation frequency ranging between 46.7% to 100% in the top 20 genes sampled by at least 3 of the 4 patients **(Supplementary Fig. 1)**. Nonsense mutations were found only in NCOR1 and TP53, with mutation frequency of 20% and 23% respectively **(Supplementary Fig. 1)**. These mutation finding highlights the genetic heterogeneity of SS and points to NCOR1, TRRAP, JAK3, PLCG1, TP53 and STAT3 (the SAAs) as initial potential targets for T-cell based immunotherapeutic intervention, based on their high mutation rate and their HLA-I peptide presentation by at least three of the four patients. This is also substantiated by their role in cancer in previous studies, that include; NCOR1 (Abedin et al., 2009; Fozzatti et al., 2013; H. Kim & Park, 2021; Perreault et al., 2006; Shu et al., 2022; St-Jean et al., 2021; Tan et al., 2019), TRRAP(Detilleux & Raynaud, 2022; Kang et al., 2018, 2023; Kwan et al., 2020; Wang et al., 2016), JAK3 (Jiang & Li, 2022; Jin et al., 2019; S. D. Li et al., 2017; Van Allen et al., 2015; Wu et al., 2023; Zhou et al., 2022),(K. B. Kim et al., 2020; F. Li et al., 2023), TP53 (Canale et al., 2022; Chen et al., 2023; B. Liu & Yi, 2022; J. Liu & Gao, 2023; Michikawa et al., 2022; Roeper et al., 2022; Sun et al., 2023) and STAT3 (Abdelhamid et al., 2022; Allam et al., 2021; Dong et al., 2021; Jin et al., 2019; Zhou et al., 2022).

Furthermore, the immunogenicity of these SAAs (NCOR1, TRRAP, JAK3, PLCG1, TP53 and STAT3) was allele-dependent and was generally highest for HLA-A*11:01 and HLA-A*02:01 and lowest for HLA-A*01:01 **(Fig. 7)**. The immunogenicity of SAAs for all the alleles was comparable, with TRRAP showing highest number of affinity peptides. These findings suggest that specific alleles may play a crucial role in shaping the immune response to SAAs in SS, with implications to personalized immunotherapy approaches. In reference to the gene expression profile of the SAAs in normal human tissues, the expression was low, except for STAT3 in select tissues **(Fig. 8)**. Thus targeting STAT3 in SS immunotherapy may lead off target, in select normal human tissues. In regards to the protein interaction partners, all SAAs had high numbers of protein interaction partners, underscoring their vital roles in SS **(Fig. 9)**. With only JAK3 and STAT3 interacting with each other at primary level, warranting future studies to determine the role of this interaction in SS.

Overall, this study finding primarily contribute to our understanding of the peptidomic landscape in SS and highlights SAAs (NCOR1, TRRAP, JAK3, PLCG1, TP53) as potential immunotherapeutic targets. The immunotherapeutic potential is based on the SAAs high mutation rate in SS, HLA-I peptides presentation, high immunogenicity, low gene expression profiles in normal human tissue, and high number of protein interaction partners. Including their role in cancer, in previous studies-mentioned above. STAT3 falls short in SAAs list as it also expressed highly in some normal human tissue, though its interaction with JAK3 at primary level, warrants future studies to determine the role of this interaction.

## Acknowledgments

The study was supported with funds from the German Ministry for Research and Education (BMBF, 13N9197), Berlin Cancer Society (WAFF200824) and the German Academic Exchange Service (DAAD). Many thanks to Dr. med. Christoph Gille, at Institute of Biochemistry, Charité-Universitätsmedizin Berlin, for his support in Sub-cellular locations and molecular functions of source proteins-data analysis.

## Author contributions

L.W.N conceived the project, conceived, designed and performed the experiments, analyzed and interpreted results, and wrote the paper. K.T-T contributed to generation of LC-MS/MS data. P.W conceived the project, contributed reagents/materials, contributed to the interpretation of the results. All authors reviewed the results and approved the manuscript.

## Figure legends

**Supplementary Figure 1:** SAAs somatic mutation types-as per catalogue of Somatic mutations in Cancer (COSMIC).

