## Supplementary figures and images for "The natural HLA-Peptidome of Sezary Syndrome: uncovering antigens for T cell-based immunotherapy"

### Supplementary Fig 1

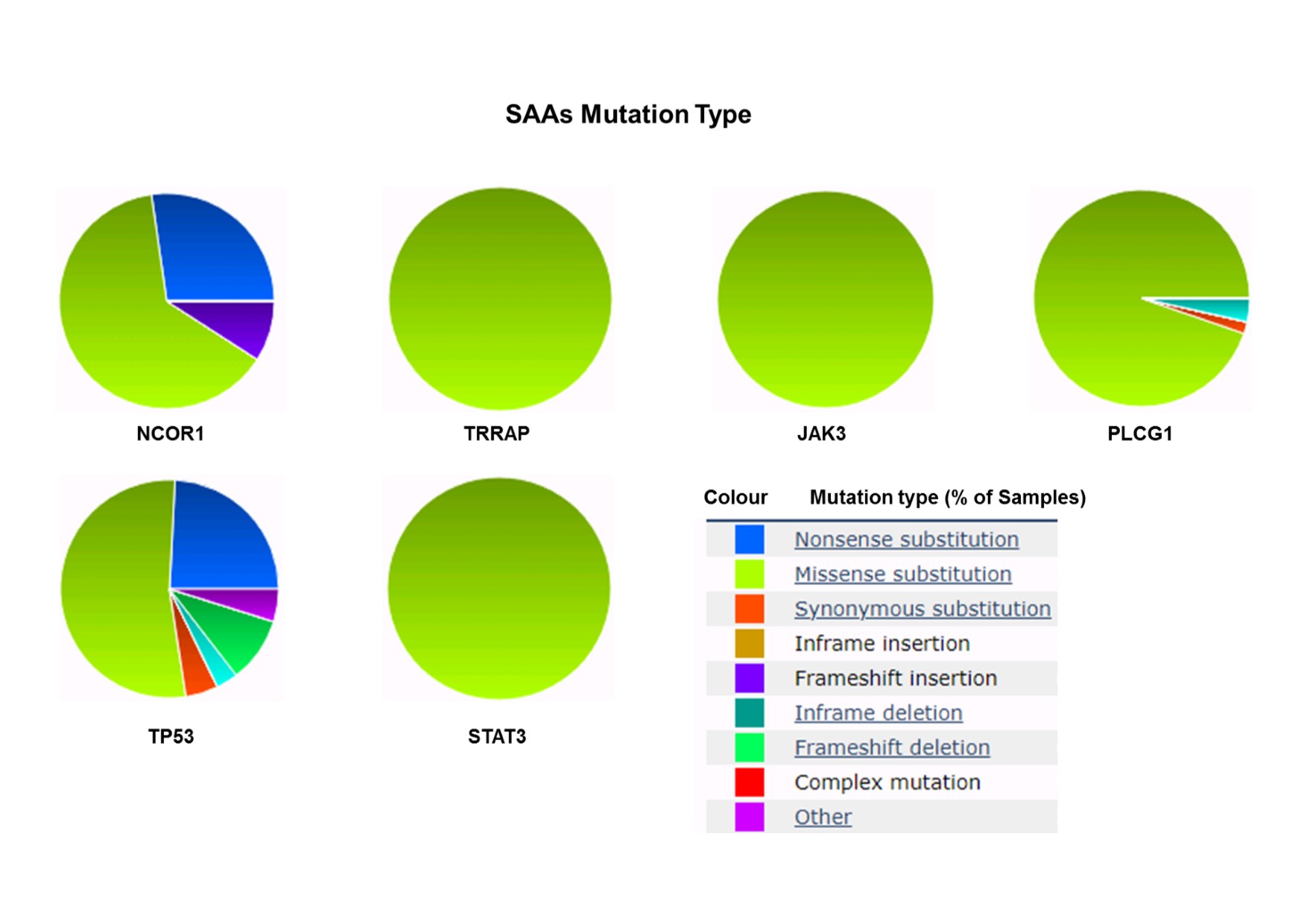
